# INSISTC: Incorporating Network Structure Information for Single-Cell Type Classification

**DOI:** 10.1101/2022.05.17.492304

**Authors:** Hansi Zheng, Saidi Wang, Xiaoman Li, Haiyan Hu

## Abstract

**Motivation:** Uncovering gene regulatory mechanisms in individual cells can provide insight into cell heterogeneity and function. Recent accumulated single-cell RNA sequencing data have made it possible to analyze gene regulation at single-cell resolution. On the other hand, understanding cell-type-specific gene regulation can also assist in more accurate cell type and state identification. Computational approaches utilizing gene regulatory relationships for single-cell type classification are under development. Methods pioneering in integrating gene regulatory mechanism discovery with cell-type classification encounter challenges such as how to accurately determine gene regulatory relationships and how to incorporate gene regulatory network structure into consideration.

**Results:** We developed a computational method to incorporate gene regulatory network structure information for single-cell type classification (INSISTC). INSISTC is capable of identifying cell-type-specific gene regulatory mechanisms while performing single cell type classification. Tested on three mouse scRNA-Seq datasets, including thousands of single-cell samples, INSISTC demonstrated its accuracy in cell type classification and its potential for providing insight into molecular mechanisms specific to individual cells. In comparison with the alternative methods, INSISTC demonstrated its complementary performance for gene regulation interpretation.

**Availability:** https://hulab.ucf.edu/research/projects/INSISTC/

**Supplementary information:** Supplementary data are available at xxxxxx online.

## 1 Introduction

Understanding gene regulatory mechanism in a cell-specific manner is a fundamental task in molecular biology. Genes are regulated at different stages, such as transcriptional and post-transcriptional gene regulation. During gene transcriptional regulation, transcription factors (TFs) and their cofactors interact with the DNA regulatory elements to regulate the gene expression levels of their target genes. Many algorithms have been developed to identify gene regulatory mechanisms through TF-target finding (Duren, et al., 2017; Sikora-Wohlfeld, et al., 2013). Many public resources have been available to store TF-target information (Fornes, et al., 2020; Han, et al., 2018; Zheng, et al., 2015).

Rapidly advanced Single-cell RNA sequencing (scRNA-Seq) enables genome-wide gene expression measurements in individual cells. scRNA-Seq data has numerous applications and has been utilized to study complicated biological processes at the single-cell resolution. For example, studying the transcriptional similarities and differences using scRNA-Seq data revealed cell-to-cell gene expression heterogeneity across species and tissues (Cao, et al., 2017; Chen, et al., 2016; Grun, et al., 2015; Rosenberg, et al., 2018). The recently accumulated scRNA-Seq-based transcriptomics data also create opportunities to understand transcriptional gene regulation at the single-cell level (Chen, et al., 2019). Computational methods have been developed to identify gene regulatory networks (GRNs), and some of them arein the context of single-cell transcriptomics (Chan, et al., 2017; Ding, et al., 2013; Matsumoto, et al., 2017; Van de Sande, et al., 2020; Wang, et al., 2017). Besides, unsupervised methods such as clustering have become common to discover cell types and cell states from scRNA-seq experiments in heterogeneous tissues (Grun, et al., 2015; Klein, et al., 2015; Zeisel, et al., 2015). Many clustering algorithms have been developed for cell-type classification using scRNA-Seq data (Guo, et al., 2015; Hao, et al., 2021; Jiang, et al., 2016; Kiselev, et al., 2017; Xu and Su, 2015; Zurauskiene and Yau, 2016). For example, Seurat V3 uses a graph clustering approach. This method projects single cells into a graph structure. Graph partitioning algorithms are then used to identify clusters. GiniClust aims to use the Gini index to identify rare cell types from scRNA-Seq data. Although these clustering algorithms have shown their capability in detecting cell types from scRNA-Seq data, they are often challenged by the lack of consistency with each other (Kiselev, et al., 2019). SC3 attempted to conquer this challenge through consensus identification. To do that, SC3 combines multiple clustering solutions to derive a consensus matrix indicating whether two cells are in one cluster. Hierarchical clustering is further applied to this matrix to obtain the final clusters.

Classification-based machine learning algorithms have also been proposed for single-cell type classification based on scRNA-Seq data (Abdelaal, et al., 2019; Alquicira-Hernandez, et al., 2019; Ma and Pellegrini, 2020; Wang, et al., 2021). These classification algorithms apply various machine learning methods such as Support Vector Machine (SVM), Random Forest, and deep learning. For example, scPred first applies a singular value decomposition-based dimension reduction approach to obtain low-dimensional principle component representations for gene expression levels (Alquicira-Hernandez, et al., 2019). Specific feature selection criteria are then applied to select informative principle components for further SVM model training and prediction. ACTINN is a recent example of deep learning-based methods for single-cell type classification (Ma and Pellegrini, 2020). ACTINN trained its deep neural network model using the Tabula Muris Atlas (a mouse cell type atlas) and a human immune cell dataset. The prediction capability was demonstrated using immune-related cell types such as mouse leukocytes and human T cell subtypes. These classification-based methods usually require training samples and specific feature selection protocols. Most of these methods utilize only the scRNA-Seq measurements of individual genes’ expression levels without considering the underlying cellular mechanisms.

Recently, another computational method called SCENIC was developed. SCENIC aims to infer single-cell-resolution gene regulatory information from scRNA-Seq data and then use this information for cell-type classification (Van de Sande, et al., 2020). SCENIC uses GENIE3 to infer TF-target co-expression relationships and uses the motif finding algorithm named RcisTarget to determine direct TF target genes. A TF and its identified targets together are defined as a regulon. SCENIC then uses AUCell algorithm to score regulon activities based on the gene expression measurements in individual cells. Using gene regulatory information to classify cell types is beneficial in two aspects. One is that integrating regulatory information is likely to help the cell type and state discovery. This is because the sensitivity of scRNA-seq technology can result in transcriptional noise (Park, et al., 2018). Also, low-expression genes are difficult to detect, causing dropouts in the data (Papalexi and Satija, 2018). The other is that cell types inferred from the underlying regulatory states can also provide insight into the cell-type-specificity of gene regulatory mechanisms. However, gene regulatory relationships forming GRNs are complex such that one TF can have many targets, and multiple TFs can collaboratively regulate the same target genes. The network structure properties in a GRN have not been taken into account for single-cell data analysis.

We developed a method called INSISTC to incorporate network structure for single-cell type classification. INSISTC utilizes the SIOMICS approach to generate a GRN with its TF-target relationships identified through de novo DNA regulatory motif discovery (Ding, et al., 2015; Ding, et al., 2014). SIOMICS is capable of considering both TFs and their cofactors for motif prediction and has demonstrated good performance. Besides, to take the structural properties of the GRN, INSISTC adopts a random-walk-based graph algorithm to represent the GRN structural information. INSISTC incorporates genes and GRN structural information by creating a Latent Dirichlet Allocation (LDA)-based topic model. The model generates cell-type-specific topics used for cell-type classification and regulatory mechanism discovery. We compared our method to SCENIC and alternative topic model construction of INSISTC. We show that INSISTC can accurately perform cell-type classification for single cells. We also demonstrated that INSISTC could uncover cell-specific gene regulatory mechanisms.

## 2 Methods

### 2.1 Overview of INSISTC

INSISTC is a topic-model-based computational framework developed to detect single cell types from scRNA-Seq measurements while providing insight into cell-type-specific gene regulatory mechanisms. For a given scRNA-Seq dataset, INSISTC consists of four steps (Fig 1). First, INSISTC provides data pre-processing and filtering. Second, based on GRN generated by SIOMICS, INSISTC executes a random walk-based graph algorithm to generate word representation for all scRNA-Seq samples. Third, INSISTC applies LDA topic model to generate a topic representation for each scRNA-Seq sample. Fourth, INSISTC performs single-cell clustering and visualization, where the SC3 and UMAP algorithms can be applied. In the following, we describe each of these four steps in more detail.

**Figure 1.**
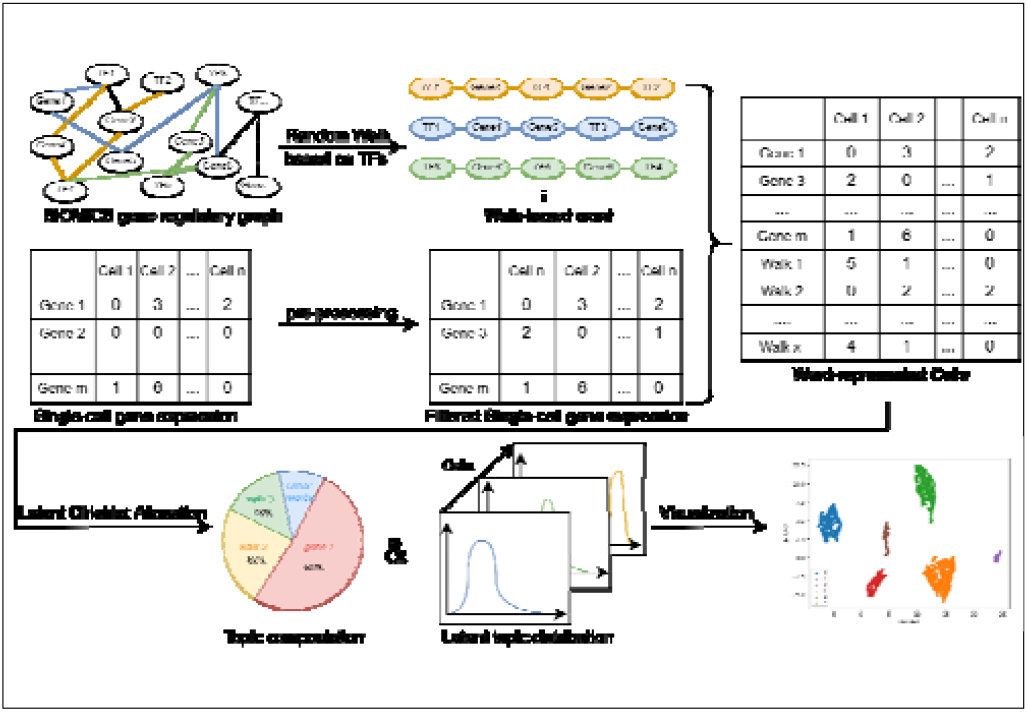
Overview of INSISTC.

### 2.2 Data collection and pre-processing

To evaluate the performance of INSISTC, we obtained three scRNA-Seq datasets: the mouse cerebral cortex data (GSE60361) (Biase, et al., 2014), the mouse skeletal muscle data (GSE143437: GSM4259473, GSM4259476, GSM4259478, GSM4259481) (De Micheli, et al., 2020), and the mouse embryo data (Array express E-MTAB-3321) (Goolam, et al., 2016). The corresponding cell numbers are 3,005, 14,242 and 124. The mouse cortex data is annotated to seven cell types, including Interneuron, pyramidal SS, pyramidal CA1, oligodendrocytes, endothelial, astrocytes ependymal, and microglia cells. The skeletal muscle data are annotated with the following 12 cell types: Mature skeletal muscle, B/T/NK cells, MuSCs and progenitors, Monocytes/Macrophages/Platelets, Endothelial, Fibro-adipogenic progenitors (FAPs), Anti-inflammatory macrophages, Resident Macrophages/APCs, Pro-inflammatory macrophages, Neural/Glial/Schwann cells, Tenocytes, and Smooth muscle cells. The mouse embryo data are annotated with five cell stages, including 2-cell stage, 4-cell stage, 8-cell stage, 16-cell stage, and 32-cell stage cells.

To filter the most likely unreliable genes that provide only noise, we applied two layers of filter on the expression matrix. The first filter is based on the total number of sequencing reads per gene. If a gene does not fit the following requirement, we remove the column of this gene from the expression matrix. The thresholds of this filter were based on each scRNA-Seq dataset, calculated by Eq. (1). If a gene with a total number of reads is less than the threshold, it will be removed from the dataset. To remove genes only expressed in one or very few cells, we applied the second filter such that genes detected in at least one percent of the total cells were kept.

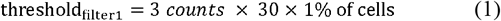

### 2.3 Computational identification of TF-target relationships

INSISTC leverages the relationship between the TFs and their target genes, constructs the gene relation graph, and traverses along the edges to infer the gene-gene connection. INSISTC uses SIOMICS v3 to obtain gene regulatory relationships. SIOMICS (systematic identification of motifs in Chip-seq data) is a computational tool for de novo discovery of motifs and TFBSs in a set of DNA sequences such as those from all peak regions of a ChIP-seq experiment. SIOMICS simultaneously considers motifs of a TF and those of its cofactors to discover motifs, which enable it to discover combinations of any number of co-occurring motifs and significantly reduce false-positive predictions compared with tools considering individual motifs separately. We call the significant motif combinations output from SIOMICS motif modules, which describes the binding pattern of a group of TFs and cofactors that co-regulate their target genes under the corresponding experimental conditions. We construct the gene regulatory network with the motif modules predicted by SIOMICS. The TF-target gene pairs for each motif were obtained by comparing the predicted motifs in motif modules with the known motifs in the JASPAR2020 vertebrate database using the tool STAMP with an E-value cutoff of 1E-5. In this way, we obtained 430 unique TFs and 20,006 unique targets, corresponding to a GRN where TFs and genes are identified as nodes, and TFs and their targets are connected by edges.

### 2.4 LDA topic model of scRNA-Seq data

INSISTC uses the LDA topic model to model a scRNA-Seq dataset. LDA is a generative probabilistic model commonly used for topic modeling (Blei, et al., 2003). LDA is motivated by the need to model a collection of discrete data. When applied to text corpora, LDA represents a document as a collection of words, and the whole word collection is defined as word vocabulary. A document can then be modeled as a finite mixture over an underlying set of topics, and a topic can be modeled as a finite mixture over an underlying set of words.

To apply the LDA to model the collections of individual cells in a scRNA-Seq dataset, we need to define the corresponding words and documents. Intuitively, we can consider the single-cell sample as a document with each gene as a word. However, defining words based on genes alone does not consider gene regulatory relationships. To account for the gene regulatory relationship, we can define each TF-target pair as a word. Nevertheless, this definition ignores the interactions between different TFs and their target genes, i.e., the structural properties o**f** a GRN.

To incorporate the structural properties of GRNs properly into the word definition of the LDA model, INSISTC uses an anchor-based random walk with a forest fire mechanism. (Hamilton, et al., 2017; Pearson, 1905). Briefly, each TF serves as an anchor for the beginning of a random walk, and each anchor is subject to a maximum of five walks. For every step of the random walk, the edge that connects the nodes of the current step with the nodes of the following step is removed to avoid a redundant walk path. The forest fire method provides for a more thorough traversal than a standard random walk, as well as a more accurate representation of the graph structure and the retention of only the unique random walk result. Each obtained random walk path is then defined a**s** a word, named as a walk-based word. The collection of all the genes and walk-based words is designated as the INSISTC vocabulary.

To further describe a single cell sample as a document with the above word definitions, we need to specify the occurrence of a specific word. INSISTC measures the occurrence of a gene based on its expression level and defines the occurrence of a walk-based word using the AUCell scoring schema (Aibar, et al., 2017). Briefly, for all the genes in a walk-based word, AUCell uses the “Area Under the Curve” (AUC) to calculate whether a critical subset of the input gene set is enriched within the expressed genes for a given single cell sample. The AUCell scores are further scaled by a constant coefficient of 100 to represent the word occurrences.

### 2.5 Comparison with SCENIC and alternative approaches

To evaluate INSISTC performance, we compared INSISTC with a recent popular method, SCENIC. We also compared INSISTC results based on alternative word and vocabulary definitions. We introduce alternative definitions including “gene-only”, “walk-only”, and “TF-target-based” vocabulary. The “gene-only” and “walk-only” are straight-forward, meaning the vocabulary only contains genes and walk-based words. To define TF-target-based vocabulary, we first filtered the TF-target gene pairs based on both genes’ expression levels; TF-target gene pairs were considered words if and only if both genes’ expression levels in the scRNA-Seq expression matrix were non-zero. For TF-target-based vocabulary, the word occurrence is the average expression between TF and target genes. The occurrence of a word for other definitions is the same as described in the above section.

To evaluate the cell type classification accuracy between any two given methods, we compare the results from different approaches with the cell type annotation from the reference publications using the adjusted rand index (ARI) (Steinley, 2004). The Rand Index (RI) can measure the similarity of two clustering results by considering the different ways of their assignments of objects to clusters. The ARI is the corrected-for-chance version of the RI. The ARI score is close to 0 if the clustering results are in a random agreement and close to 1 when the clustering results are nearly identical. The ARI is calculated based on the following equation, where denotes the number of objects in common between two clusters, and denote the sum of elements for each model.

## 3 Results

### 3.1 INSISTC reliably classifies different cell types in comparison with alternative methods

To evaluate INSISTC in terms of cell type classification accuracy, we run INSISTC on three datasets with previously annotated cell types, including mouse cortex, mouse skeletal muscle, and mouse embryo datasets (See “Materials and methods” section). The vocabulary of the topic model involved in INSISTC was defined as the union of both genes and walks (gene-walk-based vocabulary). For example, 13,063 gene-based words and 1,982 walk-based words constitute the 15,045-word vocabulary for the mouse cortex data. Similarly, for the mouse skeletal muscle data, there are 13,701 genes-based words and 2,035 walk-based words leading to 15,736-word vocabulary. The INSISTC model output major topics covered by the input single-cell samples for a specified vocabulary and topic number. Each topic contains a mixture of words that are either genes or walks. INSISTC represents each single cell sample as a mixture of the topics. Take the mouse cortex data as one example. We found 1,223 walk-based words and 1,560 gene-based words with the mixture proportion cutoff p > 0.0005. We observed that 2,886 out of 3,005 cells have at least one topic with p > 25%, 1,238 cells have at least one topic with p> 50%, and 196 cells have at least one topic with p > 75%. Of the 45 topics, 40 have at least one cell with p > 25%, 34 have at least one cell with p > 50%, and 21 have at least one cell with p > 75%. A clustering algorithm was then applied to the topics-represented single-cell samples to obtain cell type classification.

We performed SC3 clustering on topic-represented single cells to understand INSISTC results in terms of cell-type classification. SC3 is a supervised clustering tool that utilizes a consensus strategy to combine multiple clustering solutions for single-cell samples. Specification of the number of clusters is not required. To further evaluate the cell classification accuracy, we define true positives as pairs of cells with the same annotated cell type and fall into the same SC3 cluster. True negatives are cell pairs with different cell type annotations and fall into different SC3 clusters. Similarly, false negatives are cell pairs with the same cell type annotations but fall into different SC3 clusters. False positives have different cell type annotations but fall into the same cluster.

We run INSISTC under various settings of topic numbers ranging from 15 to 50 and found incorporating walk-based words in INSISTC topic discovery, in general, provides sufficient cell type classification accuracy. For the mouse cortex, skeletal muscle, and embryo samples, the best ARI achieved based on INSISTC results is 0.83, 0.67, and 0.77, respectively. In contrast, the best ARI achieved running SC3 directly on the original scRNA-Seq samples is 0.49, 0.31 and 0.58, respectively. Table 2 shows the INSISTC performance on mouse cortex data. The average recall, specificity and F1 scores are 0.73, 0.91 and 0.65. In contrast, the SC3 clustering based on the original scRNA-Seq data has the corresponding sensitivity, specificity and F1 scores as 0.28, 0.99 and 0.43 (Table 1 & supplementary Table 1). Therefore, applying the walk-incorporated topic model has a measurable impact on the ability to cluster cells belonging to the same cell types.

**Table 1.** The performance of INSISTC on mouse cortex data

| Topic num | Sensitivity | Specificity | F1-score | Cluster num | ARI |
| --- | --- | --- | --- | --- | --- |
| 15 | 0.90452 | 0.64029 | 0.55778 | 4 | 0.37418 |
| 20 | 0.83717 | 0.93290 | 0.80231 | 6 | 0.74606 |
| 25 | 0.88362 | 0.93989 | 0.83861 | 6 | 0.79217 |
| 30 | 0.77198 | 0.95326 | 0.79342 | 7 | 0.73952 |
| 35 | 0.72438 | 0.94701 | 0.75393 | 7 | 0.69080 |
| 40 | 0.58524 | 0.96586 | 0.68357 | 9 | 0.79773 |
| 45 | 0.57523 | 0.96536 | 0.67508 | 10 | 0.81575 |
| 50 | 0.53188 | 0.97448 | 0.65387 | 11 | 0.83474 |
| reference | 0.27795 | 0.99441 | 0.42803 | 7 | 0.49000 |

We also compared INSISTC results with SCENIC in terms of cell-type classification. SCENIC has recently demonstrated its capability to successfully uncover gene regulatory information and classify cell states from scRNA-Seq data. Both INSISTC and SCENIC can identify gene regulatory mechanism from single-cell transcriptomic data. INSISTC differs from SCENIC in two major aspects. One is that INSISTC focuses on network-structure incorporation using graph algorithms. The other is that in terms of regulatory motif finding, INSISTC uses SIOMICS to identify regulatory motifs that take into account binding cofactors, while SCENIC utilizes computationally defined TF-targeting relationships called regulons. We ran SCENIC (version 0.11.2) using the provided TFs and cis-regulatory database from the pySCENIC tutorial (Aibar, et al., 2017; Van de Sande, et al., 2020). We obtained 422 regulons corresponding to 422 TFs and 11,897 genes. We then performed SC3 clustering based on regulon activities inferred by SCENIC. SC3 predicted 14 clusters, based on which SCENIC corresponds to an ARI value of 0.67 comparing with the seven cell type annotations. SCENIC clustering has its overall sensitivity, specificity and F1 scores as 0.44, 0.98 and 0.59, respectively. Therefore, INSISTC has better sensitivity and F1 scores while having a slight disadvantage in specificity compared to SCENIC.

### 3.2 Network structure incorporation enhances the accuracy of cell type classifications

INSISTC defines the vocabulary of its topic model as the collection of genes and walks. To investigate how alternative vocabulary definition affects cell topic discovery and cell-type classification, we specified three alternative definitions to compare INSISTC results: “gene-only”, “walk-only”, and “TF-target-based”. Briefly, “gene-only” means that only genes are considered as words for the topic model in INSISTC, and “walk-only” means that only walk-based words are considered. TF-target-based means the TF and one of its target genes form a word to define the vocabulary.

INSISTC was run under eight topic number settings ranging from 15 to 60 for the three scRNA-Seq datasets. SC3 clustering was performed on the INSISTC topic-represented single cells. ARI was used to evaluate the cell clustering consistency with the cell-type annotation in the reference paper. The ARIs corresponding to alternative and gene-walk-based vocabulary were then compared. We found the gene-walk-based vocabulary for INSISTC, in general, resulted in more accurate cell type classification than alternative vocabulary-based INSISTC versions did (Fig 2).

**Figure 2.**
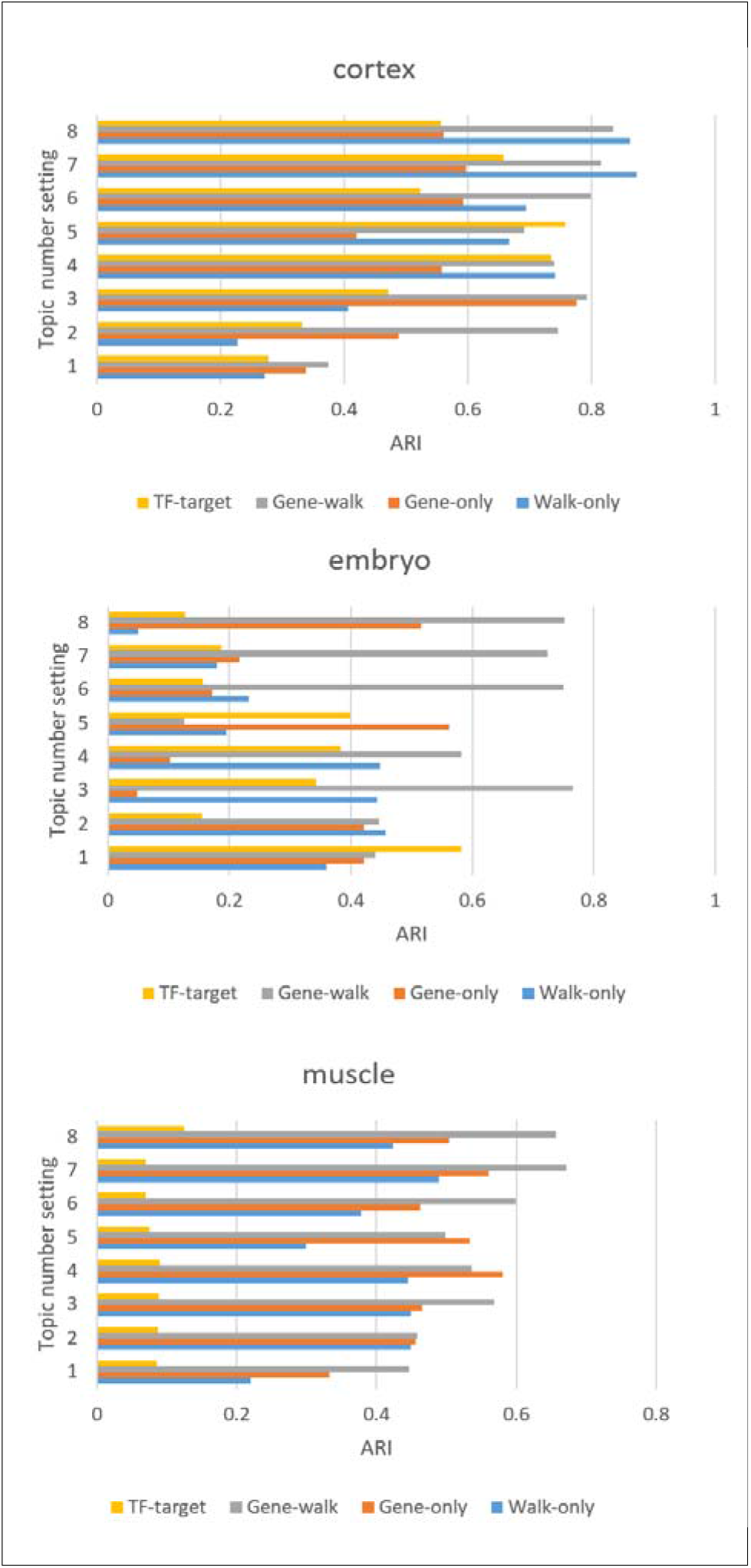
The ARIs corresponding to different vocabulary-based INSISTC results on mouse cortex, skeletal muscle and embryo datasets.

The averaged ARIs over eight topic settings are 0.72, 0.54, 0.59, and 0.54 for gene-walk, gene-only, walk-only, and TF-target-based versions. For the mouse skeletal muscle dataset, the averaged ARI over the eight topic settings are 0.55, 0.48, 0.39, and 0.09 for the same four versions. The same scenario is for the embryo dataset. The corresponding averaged ARIs are 0.57, 0.31, 0.30, and 0.29. This result shows that the network structure incorporation in the clustering procedure generally enhances the accuracy of cell type classifications.

### 3.3 INSISTC reveals marker topics contributing to cell type classification

To investigate the capability of INSISTC in interpreting the single cell type classification, we studied cell-type-specific topics (CSTs) that significantly contribute to the cell type classification. We identified CSTs based on their potential to distinguish a cell cluster from others. Using the SC3 package, we obtained CSTs as marker topics with p-values smaller than 0.01. A p-value was calculated based on Wilcoxon signed-rank test. For comparison, we also performed SC3 clustering on SCENIC results based on regulon activity scores.

For the mouse cortex dataset, we obtained 14 CSTs out of 45 topics (Fig 3). Among these topics, topic 32 can distinguish oligodendrocytes from other cell types. We found that 603 out of the 820 oligodendrocytes (74%) have topic 32 as their most enriched topic. The average enrichment proportion of the topic 32 in all the oligodendrocytes is 0.56. We performed GO analysis using 100 top-contributor words in topic 32 and found the enrichment of “response to interleukin-1” (GO:0070555, p-value: 5.12E-6) and “regulation of gliogenesis” (GO:0014013, p-value: 1.05E-5). Gliogeneisis is directly relevant to oligodendroctye generation, while Interleukin-1 has been found to regulate the proliferation and differentiation of oligodendrocytes (Vela, et al., 2002). In contrast, besides the zinc finger and BTB domain containing 33 gene (Zbtb33), SCENIC-based SC3 clustering results do not show other regulons that overlap with genes or walks in the topic 32 (Supplementary Fig. 1). Zbtb33 is involved in oligodendroglial maturation (Zhao, et al., 2016).

**Figure 3.**
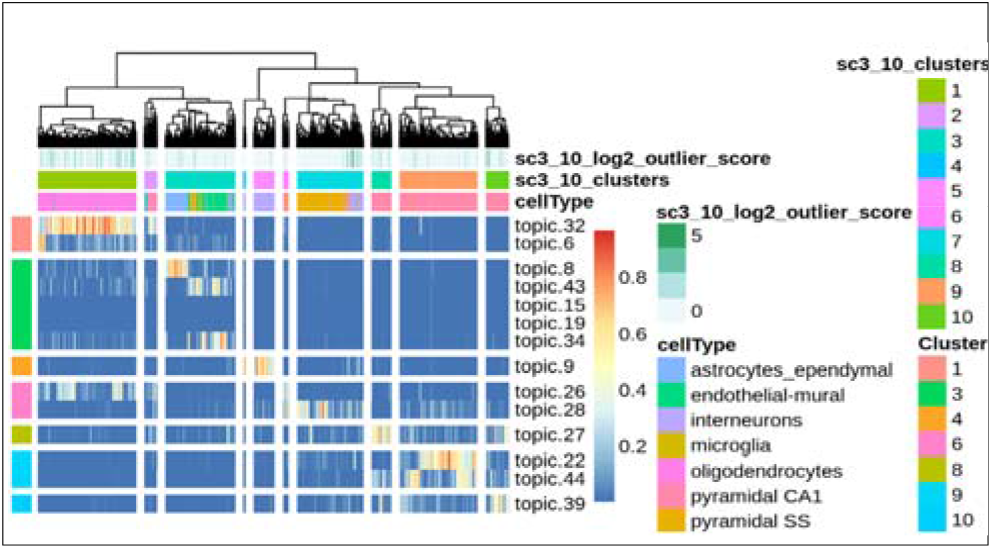
The CSTs identified in mouse cortex.

We also inspected multiple CSTs identified from the same cluster but correspond to multiple cell types. For example, topics 8, 34 and 43 were all selected as CSTs that can distinguish astrocytes and the endothelial cell type from others. Although the majority of astrocytes and endothelial cells are clustered together, topic 8 can tell astrocytes apart from others, while topic 43 is significantly enriched in endothelial cells. In both topics, Malat1 is the most enriched gene. However, a close investigation of topic 43 shows a number of top walks connected by the Kruppel-like factor 6 (Klf6) gene. This TF was reported to regulate target genes in endothelial injury recovery (Gallardo-Vara, et al., 2016).

It is interesting to see that, although topics 22, 27, 39 and 44 were all CSTs corresponding to pyramidal CA1 cell types, they actually fell into three different SC3 clusters indicating potential subtypes.

For the mouse skeletal muscle data, INSISTC identified 2,035 walk-based words and 13,071 gene-based words. Under the setting of gene-walk-based vocabulary and 55 topics, INSISTC resulted in 14 clusters corresponding to 15 cell types. We identified CSTs specific for monocytes/macrophage, endothelial, FAPs, anti-inflammatory macrophages and resident macrophages/APCs. Almost all the CSTs are supported by the GO annotation of the top-contributor words. For example, topic 14 is the CST for the anti-inflammatory macrophages. Significant GO annotation terms enriched in topic 14 include “antigen processing and presentation of peptide antigen” (GO:0048002, p-value: 1.45E-11), “immune response” (GO:0006955, p-value: 8.19E-8), “defense response” (GO:0006952, p-value: 1.65E-5), and others. Similarly, topic 1 is the CST for the FAPs. The most significant GO terms include “regulation of angiogenesis” (GO:0045765, p-value: 2.15E-5), “animal organ development” (GO: 0048513, p-value: 1.27E-5), and “positive regulation of vasculature dvelopment” (GO: 1904018, p-value: 2.35E-5).

### 3.4 INSISTC reveals cell-type-specific regulatory mechanisms

We also explored the walks in the top 100 words of CSTs to identify cell-specific regulatory mechanisms. The walk-based words ranked top according to their mixture proportions in a CST are named its top-contributor walks. We found that top-contributor walks provide insights into the regulatory mechanisms of specific cell types. Take the topic 32 identified from the cortex data for example, the GLI Family Zinc Finger 3 (Gli3) induced a number of top-contributor walks, including Rai1, Fev, Tspan2 genes. It has been shown Gli3 is important for developing mature oligodendrocytes (Tan, et al., 2006). Similarly, in topic 9, which was found to be a major marker topic for interneuron cells, we found Tcf4 involved in a few top-contributor walks that form a small network connecting Mtfr1, Dnajc4, Irf2, Foxo1, Ndufa4, Prox1, Chd1l, Pmaip1, Thy1 and others (Fig 4a). These relevant walks include TFs such as Zscan4, Hmx2, Irf2, Foxo1, and Znf16. Studies have demonstrated that Tcf4 plays an important role in the interneuron function and has been shown in interneuron dysfunction associated disorders (Jung, et al., 2018).

**Figure 4.**
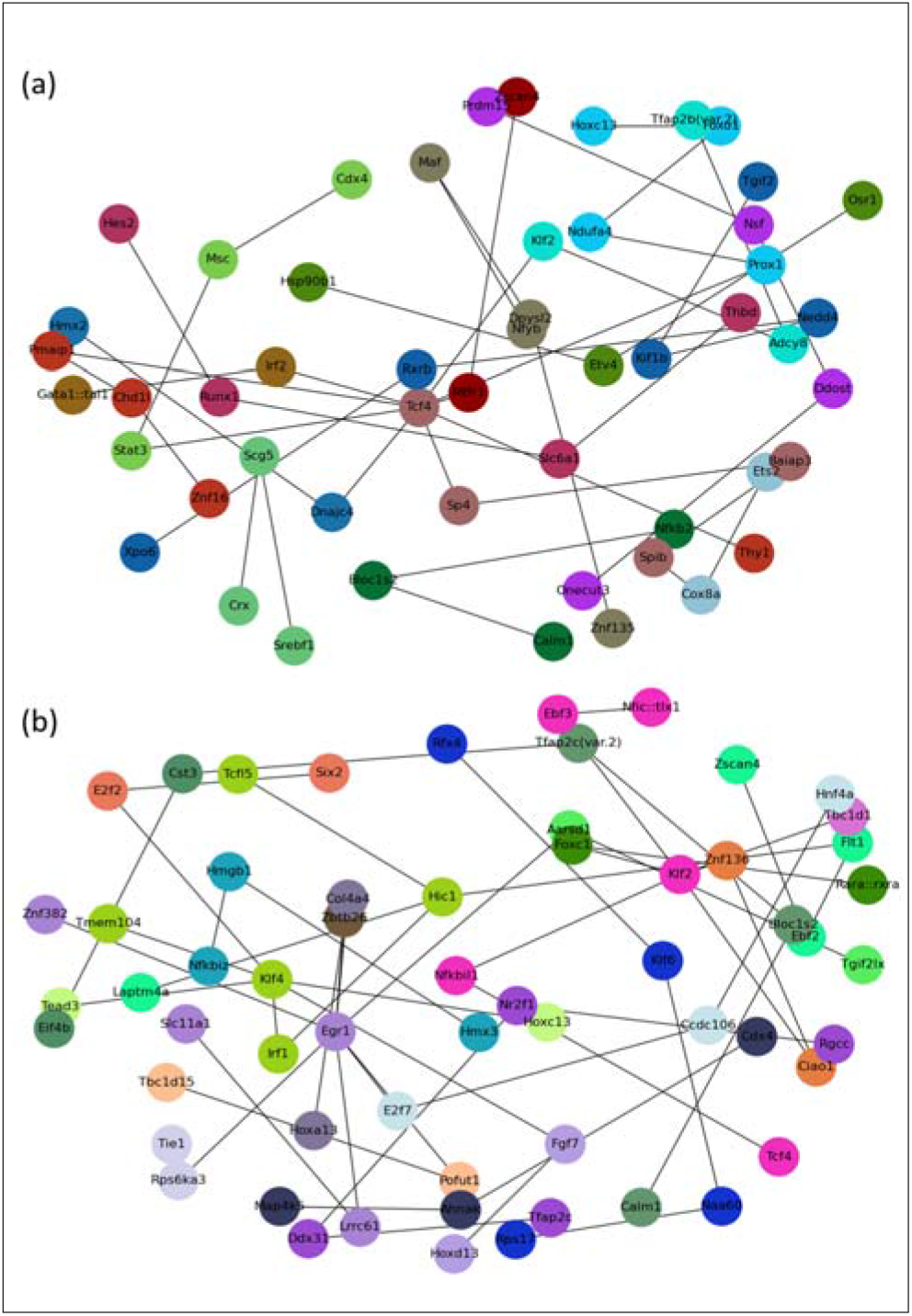
Illustration of top-contributor walks in CSTs. (a) The Tcf4 network from topic 9 in cortex. (b) the Egr1 network from topic 39 in skeletal muscle data.

For the mouse skeletal muscle data, we found the CST 39 for endothelial cells. The TF Kruppel-like factor 4 (Klf4) engaged walks were observed in the top words. Klf4 connects Tead3, E2f2, Fgf7, Irf1, Hic1, Egr2, Kif26 and other genes. Several of them, such as E2f2 and Irf1, have well-studied roles in endothelial cell growth and angiogenesis (Joyce, et al., 2004; Yan, et al., 2017). Meanwhile, Klf4 plays an important role in endothelial transcriptome regulation and greatly impacts endothelial functions (Sangwung, et al., 2017). In addition, Egr1 centered regulatory network was also revealed by the top-contributor walks involving multiple TFs such as Tgif2lx, Hoxa13, Hnf4a and Znf460 (Fig 4b). Most of these TFs participate in endothelial proliferation and angiogenesis (Shaut, et al., 2008; Yan, et al., 2017). Egr1 itself is essential to endothelial gene expression (Khachigian, et al., 1996). Similarly, the CST 14 is a marker topic for anti-inflammatory macrophages. The Irf7 and Spi1 are connected through a subnetwork that emerged from the top-container walks. Irf7 and Spi1 both play key roles in macrophage phenotype formulation and function (Gunthner and Anders, 2013; Zakrzewska, et al., 2010).

## 4 Conclusion and discussion

The availability of a large amount of scRNA-Seq data enables the study of gene regulatory mechanisms at single-cell resolution. Meanwhile, the discovery of underlying gene regulatory mechanisms can benefit more accurate cell type and state discovery from scRNA-Seq data. Methods have emerged recently to integrate gene regulatory mechanism discovery with cell-type classification. However, such method development is still at its beginning stage, and there is still space for improvement in terms of GRN construction and strategies for utilizing such GRN information. The INSISTC method was developed to overcome current challenges. INSISTC takes advantage of a de novo motif analysis that considers both TFs and their cofactors. Most importantly, INSISTC considers the graph structure of the gene regulatory network and uses a graph algorithm to incorporate this network structure. INSISTC further applies a topic model to identify particular cell-enriched topics involving cell-relevant genes and regulatory mechanisms. Such topics can be further examined for cell-specific gene regulatory mechanisms and also can be grouped, e.g., by SC3, for cell-type classification. INSISTC demonstrated sufficient cell type classification accuracy and cell-type-specific gene regulatory mechanism discovery. Compared with the recent method SCENIC, INSISTC demonstrated its complementary performance for gene regulation interpretation.

INSISTC runs SC3 clustering to identify cell types and states. This is because the SC3 algorithm offers cluster number estimation, while most clustering algorithms do not have such a function. However, any clustering algorithms can be plugged into the pipeline to derive final single-cell clusters. In addition, for the topic model that is the essential part of INSISTC, the users need to specify a topic number. There are multiple ways to determine topic numbers. For example, the topic coherence and perplexity metrics are often applied in the context of language modeling. However, it is common to run a set of topic numbers to observe biological interpretability.

INSISTC run SIOMICS to generate the TF-target relationship because SIOMICS considers both TFs and their cofactors in de novo motif discovery. However, with more accurate GRN reconstructions available, such as those integrating chromatin interaction and distal gene regulatory information (Mora, et al., 2016; Talukder, et al., 2019; Zhao, et al., 2016), the performance of INSISTC in terms of gene regulatory mechanism discovery can be further improved. Finally, although the usage of INSISTC was illustrated on GRNs, INSISTC is flexible to incorporate other types of biological networks such as pathways, protein interaction networks and gene co-expression networks. It is also possible for INSISTC to identify cell-type-specific mechanisms from a properly integrated network.

