## Supplementary for "INSISTC: Incorporating Network Structure Information for Single-Cell Type Classification"

**Supplementary Table S1**. The performance of INSISTC on mouse cortex, muscle and embryo datasets

| Sources | Topic num | sensitivity(recall) | specificity | F1 score | Cluster num | ARI |
| --- | --- | --- | --- | --- | --- | --- |
| Cortex | 15 | 0.90452 | 0.64029 | 0.55778 | 4 | 0.37418 |
|  | 20 | 0.83717 | 0.9329 | 0.80231 | 6 | 0.74606 |
|  | 25 | 0.88362 | 0.93989 | 0.83861 | 6 | 0.79217 |
|  | 30 | 0.77198 | 0.95326 | 0.79342 | 7 | 0.73952 |
|  | 35 | 0.72438 | 0.94701 | 0.75393 | 7 | 0.6908 |
|  | 40 | 0.58524 | 0.96586 | 0.68357 | 9 | 0.79773 |
|  | 45 | 0.57523 | 0.96536 | 0.67508 | 10 | 0.81575 |
|  | 50 | 0.53188 | 0.97448 | 0.65387 | 11 | 0.83474 |
|  | ref | 0.27795 | 0.99441 | 0.42803 | 7 | 0.49 |
| Muscle | 25 | 0.80492 | 0.80122 | 0.55988 | 7 | 0.44675 |
|  | 30 | 0.77197 | 0.822 | 0.56545 | 8 | 0.457678 |
|  | 35 | 0.7314 | 0.89946 | 0.64361 | 9 | 0.567656 |
|  | 40 | 0.67089 | 0.90535 | 0.61535 | 10 | 0.536633 |
|  | 45 | 0.54657 | 0.93301 | 0.57313 | 12 | 0.49818 |
|  | 50 | 0.64133 | 0.94377 | 0.65976 | 12 | 0.598557 |
|  | 55 | 0.54997 | 0.94065 | 0.58834 | 14 | 0.671519 |
|  | 60 | 0.5187 | 0.9379 | 0.55981 | 15 | 0.656894 |
|  | ref | 0.17065 | 0.99424 | 0.28401 | 63 | 0.30877 |
| Embryo | 25 | 0.62555 | 0.81869 | 0.63964 | 3 | 0.44012 |
|  | 30 | 0.62739 | 0.82192 | 0.64284 | 3 | 0.44571 |
|  | 35 | 0.97797 | 0.83803 | 0.86047 | 3 | 0.765862 |
|  | 40 | 0.64758 | 0.91539 | 0.71883 | 4 | 0.58214 |
|  | 45 | 0.54883 | 0.59488 | 0.47994 | 4 | 0.124705 |
|  | 50 | 0.88913 | 0.88235 | 0.84538 | 4 | 0.75047 |
|  | 55 | 0.86601 | 0.87712 | 0.82874 | 4 | 0.72453 |
|  | 60 | 0.8873 | 0.88477 | 0.84614 | 4 | 0.752083 |
|  | ref | 0.64758 | 0.91539 | 0.71883 | 4 | 0.58214 |

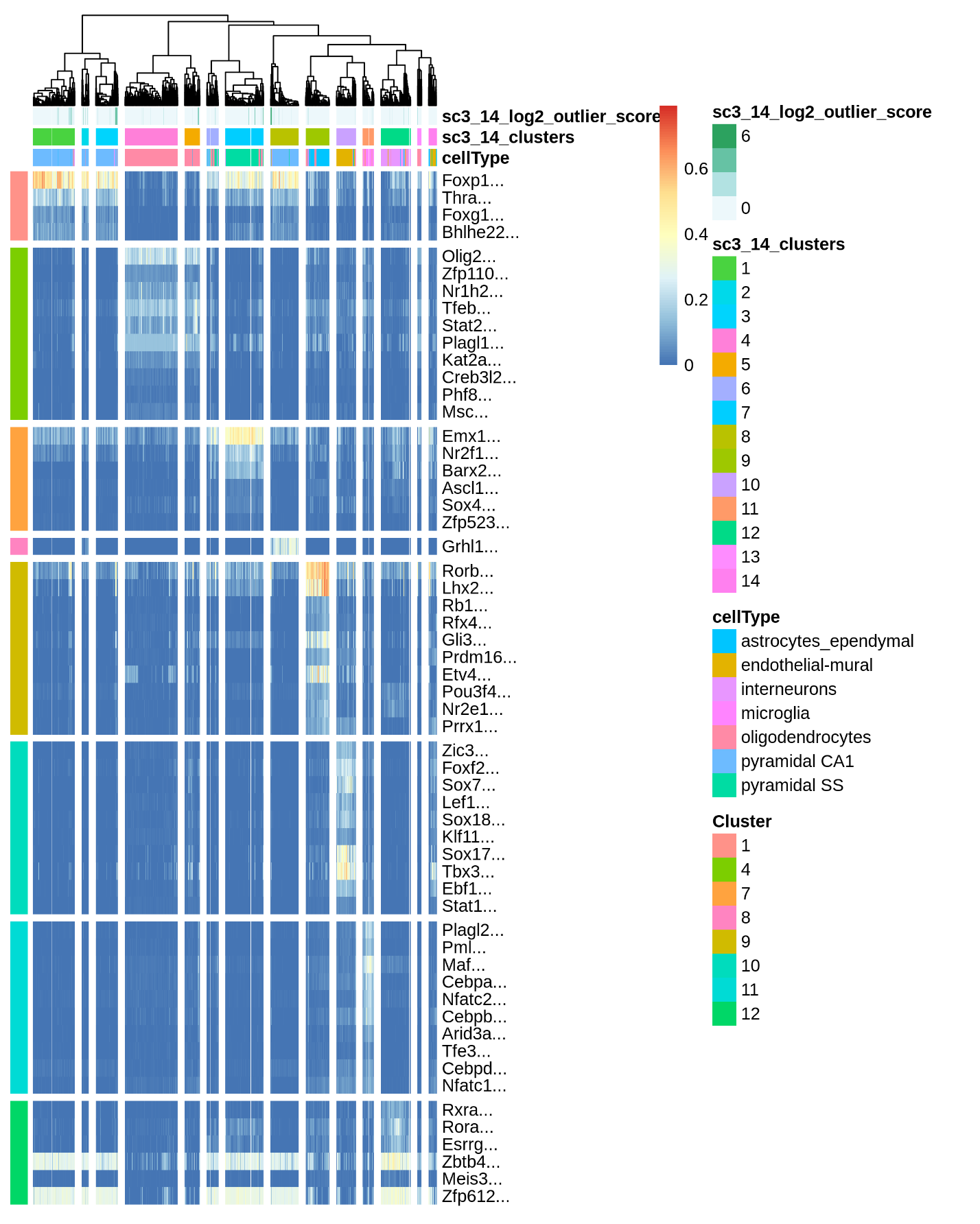

**Supplementary Fig S1**. The SCENIC-based results in mouse cortex.
